# Development of a heat-killed *fbp1* mutant strain as a therapeutic agent to treat invasive *Cryptococcus* infection

**DOI:** 10.1101/2022.12.06.519380

**Authors:** Yina Wang, Keyi Wang, Amariliz Rivera, Chaoyang Xue

## Abstract

In previous studies we determined that the F-box protein Fbp1, a subunit of the SCF(Fbp1) E3 ligase in *Cryptococcus neoformans*, is essential for fungal pathogenesis. Heat-killed *fbp1*Δ cells (HK-fbp1) can confer vaccine-induced immunity against lethal challenge with clinically important invasive fungal pathogens, e.g., *C. neoformans, C. gattii*, and *Aspergillus fumigatus*. In this study, we found that either CD4^+^ T cells or CD8^+^ T cells are sufficient to confer protection against lethal challenge of *C. neoformans* in HK-fbp1 induced-immunity. Given the potent effect of HK-fbp1 as a preventative vaccine, we further tested the potential efficacy of administering HK-fbp1 cells as a therapeutic agent for treating animals after infection. Remarkably, administration of HK-fbp1 provided robust host protection against pre-existing *C. neoformans* infection. The mice infected with wild type H99 cells and then treated with HK-fbp1 showed significant reduction of fungal CFU in the infected lung, and no dissemination of fungal cells to the brain and spleen. we find that early treatment is critical for the effective use of HK-fbp1 as a therapeutic agent. Immune analysis revealed that early treatment with HK-fbp1 cells elicited Th1 biased protective immune responses that help block fungal dissemination and promote better host protection. Our data thus suggest that HK-fbp1 is both an effective prophylactic vaccine candidate against *C. neoformans* infection in both immunocompetent and immunocompromised populations, as well as a potential novel therapeutic strategy to treat early stage cryptococcosis.

**Importance:** Invasive fungal infections, e.g., cryptococcosis, are often life threatening and difficult to treat with very limited therapeutic options. There is no vaccine available in clinical use to prevent or treat fungal infections. Our previous studies demonstrated that heat-killed *fbp1*Δ cells (HK-fbp1) in *Cryptococcus neoformans* can be harnessed to confer protection against a challenge by the virulent parental strain, even in immunocompromised animals, such as the ones lacking CD4^+^ T cells. In this study, we further determined that T cells are required for vaccine-induced protection against homologous challenge and that either CD4^+^ or CD8^+^ cells are sufficient. This finding is particularly important for the potential utility of this vaccine candidate in the context of HIV/AIDS-induced immune deficiency, the main risk factor for cryptococcosis in humans. Furthermore, in addition to the utility of HK-fbp1 as a prophylactic vaccine, we found that HK-fbp1 administration can inhibit disease dissemination when animals are treated at an early-stage during *Cryptococcus* infection. Our findings could significantly expand the utility of HK-fbp1 not only as prophylactic vaccine but also as a novel therapy against cryptococcosis. Conceptually, therapeutic administration of HK-fbp1 could have an advantage over small molecule antifungal drugs in that it is expected to have minimal side effects and lower cost. In all, our studies showed that HK-fbp1 strain can be used both preventively and therapeutically to elicit robust host protection against cryptococcosis.

## Introduction

Invasive fungal infections are emerging diseases that kill over 1.5 million people annually worldwide (1). As the immunocompromised population increases due to HIV infection, aging, and immunosuppressive treatments including transplantation and chemotherapy, etc., the incidence of invasive fungal infections is expected to rise further (1). Because fungi are eukaryotes that share much of their cellular machinery with host cells, our armamentarium of antifungal drugs is highly limited, with only three classes of antifungal drugs available (2). Among them, polyenes are toxic, triazoles are fungistatic, and echinocandins have no effect against cryptococcal infections (3). With limited drug options and the emergence of drug resistance, there is an urgent need to develop new strategies to prevent and treat invasive fungal infections to ease the public health burden they cause. For many other infectious diseases caused by viruses and bacteria, vaccines have had a transformative impact on human health and wellbeing worldwide (4). There have been numerous studies focusing on identifying fungal mutants and antigenic factors for potential fungal vaccine development and a vaccine against candidiasis has completed a phase II clinical trial (4–7). However, despite heroic efforts, there are currently no vaccines in clinical use to combat fungal infections.

*C. neoformans* is a globally distributed pathogen that causes most cases of fungal meningitis in patients with HIV/AIDS, and it is responsible for more than 180,000 deaths annually (8). People with T cell immunodeficiency, such as HIV/AIDS patients, are highly susceptible to *Cryptococcus* infection, indicating the importance of cell mediated immunity in host protection (7, 9). A wild type, *C. neoformans* strain H99 expressing IFN-γ (H99γ) has been shown to induce high Th1 immune responses and to provide full protection against virulent wild type challenge; these findings also demonstrate the importance of cell-mediated immunity (10). Indeed, *C. neoformans* mutant strains capable of inducing a highly protective Th1 response have also been reported. Several mutant strains, such as the strain overexpressing the transcription factor Znf2 (ZNF2^OE^) (11), a chitosan-deficient strain *cda1*Δ2Δ3Δ (12), and a mutant lacking sterylglucosidase (*sgl1*Δ) (13), have been identified as having increased immunogenicity in murine models. Their potential in vaccine development has also been proposed, and encouraging data have been reported (4, 7, 12, 14, 15). In addition to whole-cell vaccine strategies, simplified fungal antigenic factors, such as glucan particles, have also been identified for vaccine development (6, 16, 17). All of these exciting developments suggest that a vaccine against *Cryptococcus* or other invasive fungal infections may be feasible.

In previous studies we identified a F-box protein Fbp1, a subunit of the SCF(Fbp1) E3 ligase, and characterized the importance of the SCF(Fbp1) E3 ligase mediated proteolysis in fungal development and virulence (18, 19). We recently showed that the *fbp1*Δ mutant can trigger superior Th1 protective immunity in a CCR2^+^ monocyte-dependent manner, and found that both innate and adaptive immunity are involved in the host protection against *fbp1*Δ infection (20). The heat-killed *fbp1*Δ cells (HK-fbp1) also induce strong Th1 responses and mice vaccinated with HK-fbp1 cells show protection against challenge with the virulent parental strain (20, 21). These data indicate that the heat-killed *fbp1*Δ mutant may have potential for further development as a vaccine. Our studies also demonstrated that HK-fbp1 vaccination can trigger protection against not only its parental strain, but also against additional invasive fungal infections, including *C. neoformans, C. gattii* and *Aspergillus fumigatus*. Importantly, HK-fbp1-induced protection against *C. neoformans* is effective even in immunocompromised hosts, including animals lacking CD4^+^ T cells (21). We found that CD4^+^ T cell-depleted mice had increased CD8^+^ T cell recruitment and increased Th1 cytokine production to compensate for the loss of CD4^+^ T cells (21).

In this study, we further determined that the presence of either CD4^+^ T cells or CD8^+^ T cells is sufficient for complete protection against challenge with wild type H99 in mice previously vaccinated with HK-fbp1. Moreover, we investigated the potential therapeutic value of HK-fbp1 cells in treating infected mice. Remarkably, administration of HK-fbp1 provided robust host protection against pre-existing *C. neoformans* H99 infection. We found that early administration is critical for the therapeutic efficacy induced by HK-fbp1 cells. We determined that treatment with HK-fbp1 cells during the early stage of *Cryptococcus* infection promotes enhanced Th1 immune responses and elicits better host protection. In aggregate, our data indicate that the HK-fbp1 strain has the potential to be a suitable prophylactic vaccine candidate against invasive fungal infections, and also a potential therapeutic agent for early stage cryptococcosis.

## Materials and Methods

### Ethics statement on animal use

Female mice with an average weight of 20 - 25 g were used throughout these studies. Balb/c mice were purchased from the Jackson Laboratories, while mice of the CBA/J genetic background were purchased from Envigo. Animal studies were performed at Rutgers University Newark campus animal facility. All studies were conducted following biosafety level 2 (BSL-2) protocols and procedures approved by the Institutional Animal Care and Use Committee (IACUC) and Institutional Biosafety Committee of Rutgers University under protocol 999901066. Animal studies were compliant with all applicable provisions established by the Animal Welfare Act and the Public Health Services (PHS) Policy on the Humane Care and Use of Laboratory Animals.

### Infection with *Cryptococci*

To prepare fungal cells for infection, overnight cultures of *C. neoformans* H99 was washed three times with 1 x PBS buffer and the concentration of yeast cells was determined by hemocytometer counting. The final fungal concentration was adjusted with 1 x PBS to 2 x 10^5^ cell/ml. Each mouse was infected intranasally with 1 x 10^4^ H99 cells in a 50 μl volume after being anesthetized with a mix of Ketamine (12.5 mg/mL) and Xylazine (1 mg/mL). After infection, animals were weighed daily and monitored twice daily for progression of disease, including weight loss, gait changes, labored breathing, and fur ruffling. Over the course of the experiments, animals that appeared moribund or in pain were euthanized by CO_2_ inhalation. Survival data from the murine experiments were statistically analyzed between paired groups using the Log Rank (Mantel-Cox) test with PRISM version 8.0 (GraphPad Software) (*P* values < 0.05 were considered statistically significant). The change in body weight of each animal was calculated as follows: [(weight on day X - weight on day 0)/weight on day 0] 100%. The resulting data were plotted against time. To compare the fungal burdens, infected lungs, brains, and spleens were isolated and homogenized (Ultra-Trra T8, IKA) in 3 ml cold 1 x PBS buffer for 1 minute for each type of organ. The tissue suspensions were serially diluted and plated onto YPD agar medium with ampicillin and chloramphenicol, and colonies were counted after 3 days of incubation at 30°C.

### Vaccination strategy

*C. neoformans fbp1*Δ mutant strain was heat-killed following a previously described procedure (21). Briefly, fungal cells from YPD overnight cultures were precipitated and washed twice with sterile PBS. The cell suspension with the correct concentration was then aliquoted into Eppendorf tubes and heated on a hot plate at 75°C for 90 min. The viability of the cells following heat treatment was examined by plating the processed cell suspension on YPD agar plates; no colonies were recovered after incubation at 30°C for 3 days. Mice were vaccinated intranasally with 5 x 10_7_ heat-killed fungal cells (HK-fbp1) at day −42 unless otherwise specified. Each group of 8~10 mice were vaccinated again with the same dose of heat-killed fungal strains at day −12. A group of unvaccinated mice served as a control. The vaccinated groups and unvaccinated control group were challenged with 1 x 10^4^ live H99 cells via intranasal inoculation. Infected animals were weighed and monitored daily for disease progression, and moribund mice were euthanized. All survivors were euthanized on day 65 after challenge with live H99 cells unless otherwise specified.

### Therapeutic administration strategy

To evaluate the HK-fbp1 vaccine as a potential therapeutic agent, heat-killed *fbp1*Δ (HK-fbp1) was used to treat mice post wild type H99 challenge. The mice were infected with 1 x 10^4^ live H99 cells via intranasal inoculation at day −7 or day −3 before treatment. One group of mice were then treated intranasally with 5 x 10^7^ HK-fbp1 fungal cells at day 0. Groups of animals treated with HK-fbp1 were sacrificed at day 3 and day 7 post treatment according to the Rutgers IACUC-approved animal protocol. One group of untreated animals was sacrificed at the same time as a control. For analyzing host immune responses, bronchoalveolar lavage fluid (BALF) samples and lungs were harvested at the designated time points after treatment. Single cell suspensions of pulmonary cells were prepared for flow cytometric analysis. Infected lungs, brains, and spleens were isolated for fungal burden analysis.

### CD4^+^ / CD8^+^ T cells depletion

Mice were depleted of CD4^+^ T cell subsets or CD8^+^ T cell subsets *via* intraperitoneal administration of anti-CD4 (GK1.5, rat IgG2b) antibody (BioxCell, CAT#BE0003-1) or anti-CD8 (116-13.1, Lyt2.1) antibody (BioxCell, CAT#BE0118). Each mouse received 200 μg of GK1.5, 100 μg 116-13.1 or 200 μg isotype (LTF-2, rat IgG2b) control antibody (BioxCell, CAT# BE0090) in a volume of 200 μl PBS 9 days prior to the first vaccination, and weekly thereafter during the observation period. Efficient depletion was confirmed by measuring the prevalence of CD4^+^ or CD8^+^ T cells in blood samples by flow cytometry on the day before the first vaccination (day −43) and the day before challenge (day −1). The depletion was also confirmed by measuring the prevalence of CD4^+^ or CD8^+^ T cells in BALF and lung tissues by flow cytometry at the endpoint of the experiment. The anti-CD4 antibody used for flow cytometric analysis binds to epitope of the CD4 protein at locations distinct from GK1.5. The RM4-4 FITC Rat Anti-mouse CD4 antibody (BD Biosciences, CAT#553055) and YTS169.4 PE Rat Anti-mouse CD8 antibody (Invitrogen, CAT#MA5-17606) were used for blood samples flow cytometric analysis, while the CD4 RM4-5 BV421 Rat Anti-mouse CD4 antibody (BD Biosciences, CAT#100543) and Rat Anti-mouse CD8 antibody (Invitrogen, CAT#MA5-17606) were used for BALF and lung tissues flow cytometric analysis.

### Depletion confirmation

To confirm the efficiency of CD4^+^ or CD8^+^ T cells depletion, animal blood samples were processed for flow cytometry as described previously (21). Blood samples (2-3 drops/tail) were collected and placed into 50 μl of heparin (100 USP Heparin units/ml) in a 96-well plate. After collection, bleeding was stopped by applying Kwik Stop styptic powder using moistened cotton applicator to end of tail. Blood cells were washed with 150 μl of 1 x PBS and resuspended in 200 μl of Red Blood Lysis buffer (NH_4_Cl 155mM) and (NaHCO_3_ 10mM), pH 7.2. Cells were washed again with 200 μl 1 x PBS and resuspended in 50 μl of 1:50 dilution of Fc block (CD16/CD32, 2.4G2) in FACS buffer (0.1% sodium azide in 1 x PBS). After incubation on ice for 15-20 min, cells were washed with 150 μl of FACS buffer, and resuspended in 50 μl of the appropriate antibody mix for each strain. Following 45-60 mins co-incubation on ice, samples were washed with FACS buffer and resuspended in 200 μl of FACS buffer for flow cytometry. Cell surface antibodies CD45 (30-F11 BUV395), Thy1.2 (53-2.1 PE-Cy7), CD4 (RM4-4 FITC) CD8α (53-6.7 Pacific blue) were used to confirm CD4^+^ or CD8^+^T cells depletion.

### Lung processing

Single cell suspensions of pulmonary cells were prepared for flow cytometric analysis as previously described (22). In brief, lung tissue was minced in 5 ml of 1 x PBS containing 3 mg/ml collagenase type IV (Worthington). Samples were incubated at 37°C for 45 min and washed with 1 x PBS three times. After digestion, residual RBCs were removed using RBC lysis buffer (155 mM NH_4_Cl and 10 mM NaHCO_3_, PH 7.2). Lung cell suspensions were used for flow cytometry. Lungs single cell suspensions were stained for monocytes [CD45 (30-F11BUV395), CD11b (M1/70 PerCP Cy5.5) and Ly6C (AL-21 PE)], Mo-DCs [CD45 (30-F11 BUV395), CD11b (M1/70 PerCP Cy5.5), Ly6C (AL-21 PE), CD11c (HL3 BV510) and MHC Class II I-A/I-E (M5/11.415.2 BV711)], neutrophils [CD45 (30-F11 APC-Cy7), CD11b (M1/70 PerCP Cy5.5), Ly6C (AL-21 PE) and Ly6G (1A8 APC)], CD4 T cells [CD45 (30-F11 BUV395), CD4 (RM4-5 BV421)], CD8 T cells [CD45 (30-F11 BUV395), CD8α (53-6.7 BV711). All antibodies used for lung staining were from BD Biosciences. All samples were analyzed using BD LSRFortessa^™^ Flow Cytometer and FlowJo software (Tree Star, Inc).

### Intracellular cytokine staining of T cells harvested in BALF and Flow Cytometry

For analyzing host immune responses, bronchoalveolar lavage fluid (BALF) samples were harvested at the endpoint after inoculation. BALF was collected in 3 ml of 1 x PBS buffer using a catheter inserted into the trachea of animal post-euthanasia, and airway-infiltrating cells were lavaged with ~ 1 mL of 1 x PBS at a time to a total volume of 5 mL. RBCs were removed using RBC lysis buffer. BALF cells were then plated in a 96-well round-bottom plate and restimulated using BD-Leukocyte Activation Cocktail containing BD GolgiPlug^™^ (BD Biosciences) according to the manufacturer’s instructions. Six hours after activation, BALF cells were surface-stained with fluorescently labeled antibodies against Thy1.2, CD4 and CD8. Samples were fixed in 1% paraformaldehyde overnight. Prior to intracellular staining the samples were permeabilized with 1 x BD Perm/Wash buffer according to the manufacturer’s instructions. Intracellular cytokine staining (ICCS) was done using fluorescently labeled antibodies against IFN-γ, IL-17A, TNF-α and IL13 diluted in 1 x BD Perm/Wash for 30 minutes on ice. Samples were immediately washed and analyzed by flow cytometry as described below. BALFs were cell surface stained for T cells with Thy1.2 (53-2.1 PE-Cy7), CD4 (RM4-5 BV421) CD8α (53-6.7 BV711) and ICCS for IFN-γ (XMG1.2 PE), IL-17A (eBio17B7 APC), TNF-α (MP6-XT22 BV711) and IL-13 (eBio13A FITC) following standard procedures. Most antibodies and reagents for cell surface and ICCS were from BD Biosciences, except IL-17A and IL-13 which were obtained from eBioscience, Inc.

### CD4^+^ T cell isolation and CD4^+^ T cell recall response

Antigen-presenting cells (APCs) were prepared from the spleen of syngeneic, uninfected donor mice. Splenic cell suspensions were depleted of T cells by antibody complement-mediated lysis. Splenic cells were incubated with anti-Thy1.2 antibodies and rabbit complement (Low Tox; Cedarlane Labs) at 37 °C for 1 hour. Lung-draining lymph nodes (MLNs) were collected and placed in a 10ml of 1 x PBS. Total lymphocyte cell suspensions were prepared by gently releasing the cells into the 1 x PBS by applying pressure to the lymph nodes with the forested ends of two glass slides. Repeated pressure was applied until the tissue was reduced to the smallest size possible. Samples were collected and processed in the same way individually. For CD4 T cell isolation, individual samples from each group were pooled (5 mice). CD4^+^ T cells were purified using a negative-sorting CD4^+^ isolation kit (Miltenyi Biotec, Inc.). CD4^+^ T cell isolation was done following the manufacturer’s instructions and were consistently found to be > 90% pure, as assessed by flow cytometry. Purified CD4^+^ T cells (2×10^5^) were cultured with T cell-depleted antigen-presenting cells (APCs; 3×10^5^) in RPMI containing 10% fetal calf serum (FCS), Penicillin-Streptomycin (2200 U/ml, Gibco^™^) and Gentamicin sulfate solution (1 mg/ml). The cultures were plated in flat-bottom 96-well plates and incubated at 37°C with 5% CO_2_ for 72 hours. To measure *Cryptococcus*-specific CD4+ T cells responses, CD4-antigen-presenting cell cultures were incubated with sonicated (Qsonica Sonicator Q55) H99 yeasts as a source of fungal antigens. The amount of antigen used was adjusted to a multiplicity of infection of 1:1.5 (antigen-presenting cell:yeast ratio). The fungal growth inhibitor voriconazole was used at a final concentration of 0.5 mg/ml to prevent any fungal cell outgrowth during the culture period. After 72 hours of culture at 37 °C with 5% CO_2_, supernatants were collected for cytokine analysis by ELISA (IL-2 BD-OptEIA^™^; IL-17A, IFN-γ ThermoFisher) following manufacturer’s instructions.

## Results

### CD4^+^ or CD8^+^ T cells are sufficient for full protection against wild type H99 challenge in HK-fbp1-vaccinated mice

Individuals with immunodeficiency, such as impaired T cell function in HIV/AIDS patients, are highly susceptible to *C. neoformans* infection, suggesting the importance of T cells in defense against cryptococcosis (9). Our previous studies demonstrated that the *fbp1*Δ mutant elicited superior protective Th1 host immunity in the lungs and that the enhanced immunogenicity of heat-killed *fbp1*Δ (HK-fbp1) yeast cells can be harnessed to confer protection against a subsequent infection with the virulent parental strain in immunocompetent or CD4^+^ T cells deficient hosts (20, 21). Given the clinical significance of T cell deficiency to the susceptibility to cryptococcosis in patients, it is also critical to know whether host protection can be established following HK-fbp1 vaccination in both CD4^+^ and CD8^+^ T cell-deficient hosts. Therefore, we further examined the potential vaccine protection in mice depleted of CD4^+^ and/or CD8^+^ T cells. An animal model of T cell deficiency was achieved by administration of 200 μg/mouse anti-CD4 antibody and/or 100 μg/mouse anti-CD8 antibody 9 days prior to first vaccination, and weekly thereafter during the course of the experiment (Fig. 1A). Efficient depletion was confirmed by measuring the prevalence of CD4^+^ T cells and CD8^+^ T cells in blood samples by flow cytometry on the day before the first vaccination (day −43) and the day before challenge (day −1) (Fig. S1). We tracked the changes in animal body weight weekly throughout the experiment. We noticed that CD4^+^ T cell-depleted, CD8^+^ T cell-depleted and isotype control animals challenged with H99 cells maintained or increased in body weight over time, while the CD4^+^ and CD8^+^ T cells double depleted mice and unvaccinated ones lost weight rapidly following infection (Fig. 1B). Consistent with animal body weight changes, mice administered with either anti-CD4 antibody, anti-CD8 antibody, or isotype antibody control survived for other two months after challenged with H99. Thus, HK-fbp1-vaccinated animals depleted of CD4^+^ or CD8^+^ T cells were fully protected against H99 challenge (Fig. 1C and Fig. S1). The difference between the median survival time for vaccinated mice with CD4^+^ and CD8^+^ double T cells depletion and unvaccinated mice was not statistically different, suggesting that vaccination does not work in mice that lack both T cell subsets (Fig. 1C).

**Figure 1.**
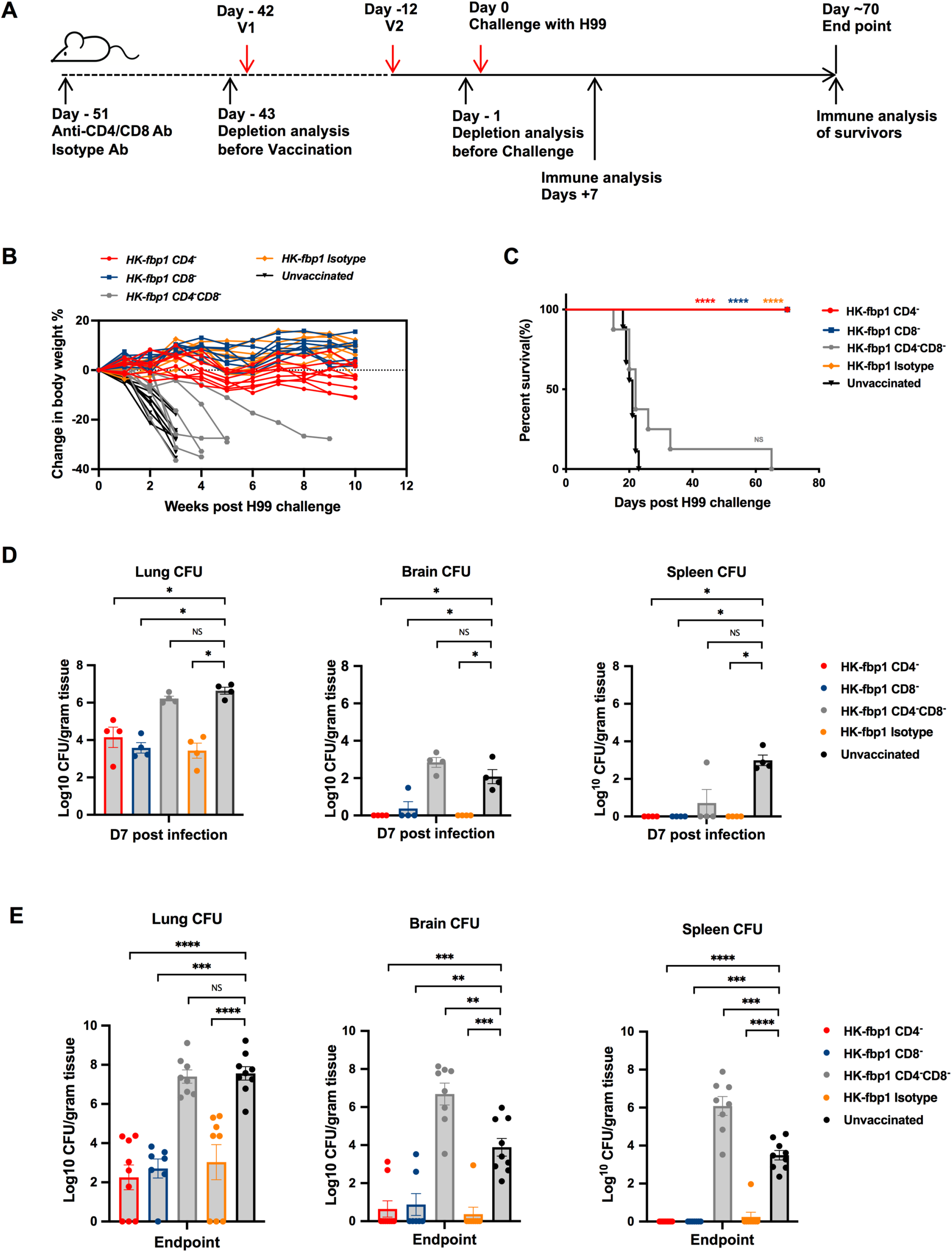
Animals with depleted either CD4^+^ T cells or CD8^+^ T cells remain fully protected by HK-fbp1 vaccination. **(A)** Strategy of vaccination. Depletion of CD4^+^ T cells and/or CD8^+^ T cells was accomplished by injecting anti-CD4 antibody (GK1.5, rat IgG2b), anti-CD8 antibody (116-13.1) weekly, starting 9 days prior to the first vaccination, and weekly thereafter during the course of the experiment. **(B)** Dynamic changes of body weight in vaccinated and unvaccinated animals after challenge with 1×10^4^ H99 cells. All live mice from each group were weighted, and their average weight changes were presented. **(C)** Survival curves of CBA/J mice vaccinated with 5×10^7^ HK*-* fbp1 and challenged by 1×10^4^H99 cells. “****”, P<0.0001 (determined by Log Rank [Mantel-Cox] test). **(D)** Fungal burden in the lungs, brains and spleens of vaccinated animals infected with 1×10^4^ H99 cells at day 7 post-infection and the fungal burden in unvaccinated control animals. Each symbol represents one mouse. Bars represent the mean ± the standard error of the mean. “*”, P<0.05 (determined by Mann-Whitney test). **(E)** Fungal burden in the lungs, brains and spleens of vaccinated animals infected with 1×10^4^ H99 cells at the end point and the fungal burden in unvaccinated control animals. Bars represent the mean ± the standard error of the mean. “****”, P<0.0001; “***”, P<0.001; “**”, P<0.01 (determined by Mann-Whitney test).

At 7 days post challenge, five mice from each group were sacrificed and examined for fungal burden in lungs, brains, and spleens. For CD4^+^ T cell-depleted or CD8^+^ T cell-depleted animal groups and the isotype control group, no CFUs were detected in all but one infected mouse brains and spleens, and significantly lower CFUs were detected in infected lungs compared to the lungs of non-vaccinated animals. The fungal burden in the CD4^+^ and CD8^+^ T cells double depleted mice is similar to the unvaccinated control group, indicating that the presence of either CD4^+^ or CD8^+^ T cells is necessary and also sufficient to induce protective immunity against *C. neoformans* challenge (Fig. 1D). We also analyzed the endpoint (day 70) fungal burdens in these animal groups and observed significantly reduced fungal burdens in the lungs of vaccinated animals, except the CD4^+^ and CD8^+^ T cells double depleted mice. Most of the brains and spleens of immunized animals were cleared of H99 cells during the period of this experiment, suggesting that immunization helped CD4^+^ or CD8^+^ T cell-deficient animals clear or restrict *Cryptococcus* cell proliferation (Fig. 1E). Altogether, our studies showed the HK-fbp1 vaccination induced protection is dependent on T cells, and the vaccine remains effective in immunocompromised hosts that lack either CD4^+^ or CD8^+^ T cells, but not both.

### Robust cytokine production by either CD4 or CD8 T cells underlies HK-fbp1-induced vaccine protection

To understand how protection is established in either CD4^+^ or CD8^+^ T cell-depleted mice, we examined the immune responses of remaining CD4^+^ or CD8^+^ T cell populations in vaccinated and unvaccinated mice at day 7 post challenge and also at the end point of the experiment (day 70). At day 7 post infection, we found that while the isotype control treated mice had a significant increase in the number of CD4^+^ T cells in the infected lungs, CD4^+^ T cells were not detectable in the BALF of CD4^+^ T cell depleted mice, and CD8^+^ T cells were not detectable in the lungs of CD8^+^ T cell depleted mice as expected (Fig. 2A). CD4^+^ or CD8^+^ T cell depletion were maintained weekly until the endpoint of the experiment (day 70). We examined T cells in lung, BALF and lung draining lymph nodes (mLN) at the endpoint, we confirmed that CD4^+^ or CD8^+^ T cells were completely depleted throughout the whole experiment (Fig. S2).

**Figure 2.**
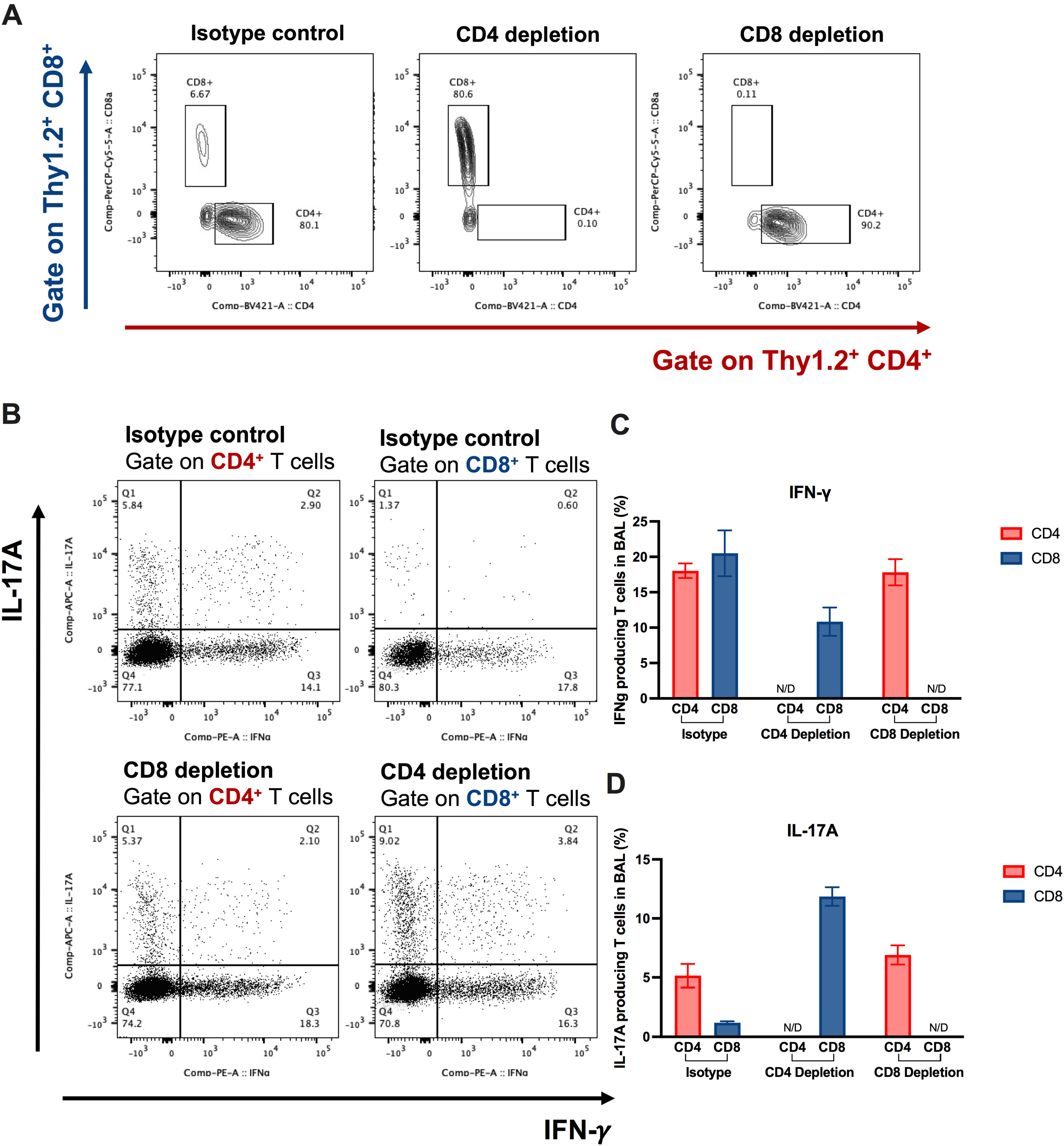
Role of CD4^+^ and CD8^+^ T cells and enhanced Th1 and Th17 T cell responses during the induction of protective immunity by HK-fbp1 in immunodeficient animal model. **(A)** Representative FACS plots of CD4^+^ T cells and CD8^+^ T cells in the BALF of isotype control mice (left), CD4^+^ T cells depleted mice (middle), and CD8^+^ T cells depleted mice (right). **(B)** Representative FACS plots of cytokine production gate on CD4^+^ T cells or CD8^+^ T cells in isotype control mice (top), CD4 depleted mice (bottom), and CD8 depleted mice (bottom) at day 7 post infection. **(C and D)** Cytokine expression analyzed by ICCS. Plots of cytokine production in CD4^+^ T cells gated as Thy1.2^+^ CD4^+^ CD8^-^ T cells. Plots of cytokine production in CD8^+^ T cells gated as Thy1.2^+^ CD4^-^ CD8^+^ T cells. The frequencies of IFN-γ **(C)**, IL-17A **(D)** producing CD4^+^ or CD8^+^ T cells in BALF were analyzed as shown in panel.

Previously, we determined that IFN-γ is a critical mediator of vaccine-induced protection in this model (20, 21). Mice defective in IFN-γ responsiveness were not protected after HK-fbp1 vaccination and were as susceptible to infection as unvaccinated mice (21). Therefore, to understand how remaining T cell populations protect vaccinated mice against challenge with H99, we examined the cytokine responses of the remaining CD4^+^ T cells or CD8^+^ T cells in the airway at day 7 post infection by intracellular cytokine staining. Our data show that CD8^+^ T cells produce IFN-γ and IL-17A in CD4^+^ T cell-depleted mice (Fig. 2). In the isotype control treated mice, these protective cytokines were mainly produced by CD4^+^ T cells (Fig. 2). Meanwhile, in the CD8^+^ T cell deleted mice, CD4^+^ T cells are robust producers of protective cytokines (Fig. 2B, C, D). These results suggest that CD8^+^ T cells are able to produce protective cytokines, which can compensate for the lack of CD4^+^ T cells. Our findings thus indicate that CD4^+^ T cells or CD8^+^ T cells are sufficient to confer protection against challenge with H99 in HK-fbp1-vaccinated mice due to their ability to produce protective IFN-γ and IL-17.

### Treatment of *C. neoformans* H99 infected mice with HK-fbp1 vaccine inhibits fungal dissemination

A therapeutic vaccine is one in which the vaccine is used after infection occurs, aiming to induce anti-infective immunity to alter the course of disease (23). It works by activating the host immune system to fight an infection. Since the HK-fbp1 vaccine candidate confers high protection against *Cryptococcus neoformans* infection prophylactically by inducing T cell-dependent protective immunity, we asked whether HK-fbp1 could be applied as a therapeutic agent to treat *Cryptococcus-infected* hosts by boosting their immunity. To test this idea, previously naïve mice were infected with 1 × 10^4^ live H99 cells via intranasal inoculation. At day 3 post infection, the infected mice were treated intranasally with 5 × 10_7_ HK-fbp1 cells. Treated and untreated animals were monitored for survival for up to 70 days (Fig. 3A).

**Figure 3.**
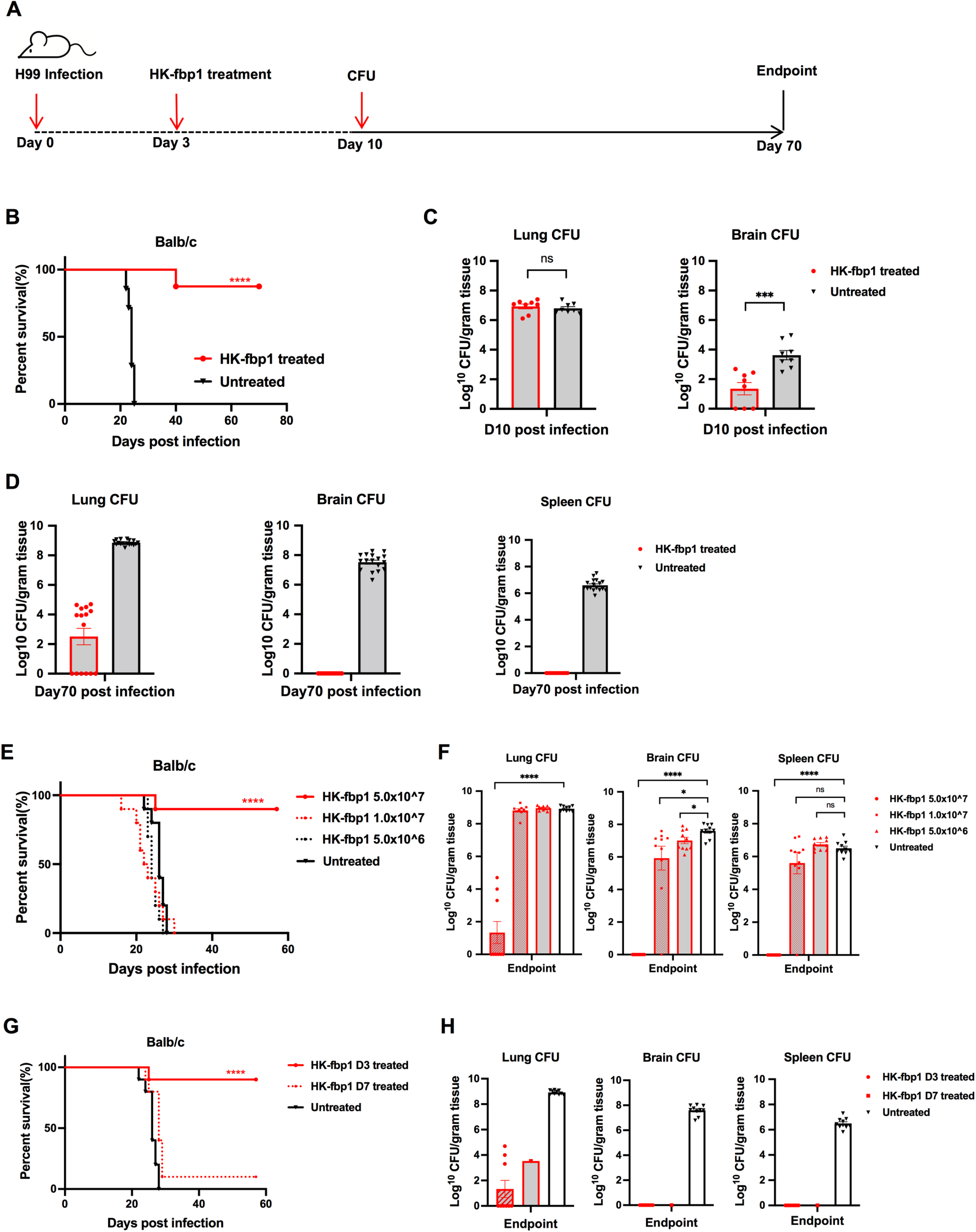
HK-fbp1 treatment of mice infected by *C. neoformans* H99 blocked fungal dissemination to cause lethal infection. **(A)** Scheme of HK-fbp1 treatment strategy. **(B)** Survival curves of HK-fbp1 treated Balb/c mice challenged with *C. neoformans* H99. “****”, P<0.0001 (determined by Log Rank [Mantel-Cox] test). **(C)** Fungal burden in the infected lungs, brains and spleens at day 7 post-treatment. Each symbol represents one mouse. Bars represent the mean ± the standard error of the mean. “***”, P<0.001 (determined by Mann-Whitney test). **(D)** Fungal burden in the infected lungs, brains and spleens at the end of the experiment (~70 days post challenge). Each symbol represents one mouse. **(E)** Survival curves of mice pre-challenged by infection with 10^4^ wild type H99 cells and treated with different doses of HK-fbp1 at day 3 post infection. 8~10 female Balb/c mice were used for each group. “****”, P<0.0001 (determined by Log Rank [Mantel-Cox] test). **(F)** Fungal burden in the lungs, brains and spleens of HK-fbp1 treated survivors at the end of the experiment and fungal burden of untreated control mice. Bars represent the mean ± the standard error of the mean. “****”, P<0.0001; “*”, P<0.05 (determined by Mann-Whitney test). **(G)** Survival curves of mice pre-challenged with 10^4^ wild type H99 cells at 3 days and 7 days before treated with high dose of HK-fbp1. “****”, P<0.0001 (determined by Log Rank [Mantel-Cox] test). **(H)** Fungal burden in the lungs, brains and spleens of the surviving mice at the end of the experiment and the fungal burden in untreated mice.

Remarkably, HK-fbp1 treatment induced significant protection against preexisting H99 infection. All untreated mice succumbed to fatal infection at ~ 20 days, while most treated animals survived for over 70 days following HK-fbp1 treatment at day 3 post infection (Fig. 3B). We examined the fungal burden in the lungs, brains and spleens of the treated and untreated mice at set timepoints of the experiment. Although no significant difference in fungal CFUs were detected between HK-fbp1 treated lungs and the untreated controls at 7 days post-treatment, significantly reduced CFUs were detected in most of the brains of treated animals (Fig. 3C). Endpoint fungal burden analysis revealed significantly reduced fungal burdens in the lungs of HK-fbp1 treated animals. Importantly, none of the surviving mice of the treated group displayed extrapulmonary dissemination of H99 cells to the brain and spleen (Fig. 3D). H99 cells were totally cleared from all brains and spleens of treated animals during the period of the experiment, indicating that HK-fbp1 treatment is sufficient to clear or restrict H99 cells from dissemination, likely by inducing protective immunity. Taken together, our results demonstrate that treatment with the HK-fbp1 vaccine protects animals from pre-existing lethal infection and restrains proliferation of wild type H99 cells in the lung.

### High dose and early treatment are required for the therapeutic efficacy induced by HK-fbp1 against *Cryptococcus* infection

Our previous studies revealed that as a prophylactic vaccine, the efficacy of HK-fbp1 vaccine protection in mice was dose dependent (21). Mice immunized with higher doses of vaccine exhibited better protection than those vaccinated with low doses (21). Mice vaccinated with 5 × 10^7^ HK-fbp1 conferred 100% protection, while no clear protection was observed for mice vaccinated with 5 × 10^6^ HK-fbp1 cells (21). Since our data showed a potential therapeutic value for our vaccine as treatment for animals with pre-existing infection, we examined the potential dose effect of the HK-fbp1 treatment. Mice were administrated intranasally with different HK-fbp1 inocula (5 × 10^7^, 1 × 10^7^, 5 × 10^6^, cells/mouse) at day 3 post infection. We did find a dose dependent efficacy (Fig. 3E). Mice treated with higher dose of HK-fbp1 exhibited better protection than those treated with low doses. Mice treated with 5 × 10^7^ HK-fbp1 conferred ~90% protection, while no clear protection was observed for mice treated with 1 × 10^7^ and 5 × 10^6^ HK-fbp1 cells (Fig. 3E). Examination of the fungal burden in high dose treated mice that survived H99 infection showed that fungal cells were cleared from the brains and spleens, while lungs had significantly less fungal loads (Fig. 3F). Overall, our data suggest that a fungal antigenic threshold has to be reached in order to induce efficient protection with HK-fbp1 treatment.

We then tested whether HK-fbp1 treatment remains effective on infected animals with disseminated disease. Since *C. neoformans* infection typically disseminates at day 7 post-infection intranasally, while it remains as a local lung infection at day 3, we set two treatment timepoints: 7 days post-infection as long-term infection and 3 days postinfection as short-term infection. We challenged mice at −7 days, −3 days before HK-fbp1 treatment (Fig. 3G). Hk-fbp1 treatment at day 3 post infection conferred ~90% protection, while no clear protection was observed for mice treated at day 7 post infection (only 10% mice survived for 70 days). Examination of the fungal burden in mice treated early showed clearance of fungal cells in the brains and spleens, and infected lungs had significantly less fungal loads (Fig. 3H). Similar results were observed for the one surviving mouse treated at day 7 post-infection (Fig. 3H). Overall, our data indicate that high dose and early treatment are required for the efficacy of the HK-fbp1 vaccine to treat *Cryptococcus* infection.

### Early treatment of HK-fbp1 induces a protective host response and prevents fungal dissemination

Our previous study demonstrated that a strong Th1 immune response developed in mice immunized with HK-fbp1 cells (20, 21). Because mice treated with HK-fbp1 at early time post-infection showed strong protection and blocked fungal dissemination, we set out to examine the immune response of *Cryptococcus*-infected mice that were treated with HK-fbp1. Mice were infected with H99 for 3 days or 7 days and then treated with HK-fbp1, respectively (Fig 4A). Our data showed that at day 10 post infection, fungal CFU was detected in both day 3-treated mice (early treatment), and day 7-treated (late treatment) mice. Although there is no difference of CFU in the lung of day 3-treated mice compared to 7-treated mice, the day 3-treated mice showed significantly reduced fungal CFU in infected brains, suggesting an inhibition of fungal dissemination, while the day 7-treated mice showed a similar fungal burden as those untreated mice (Fig. 4B).

**Figure 4.**
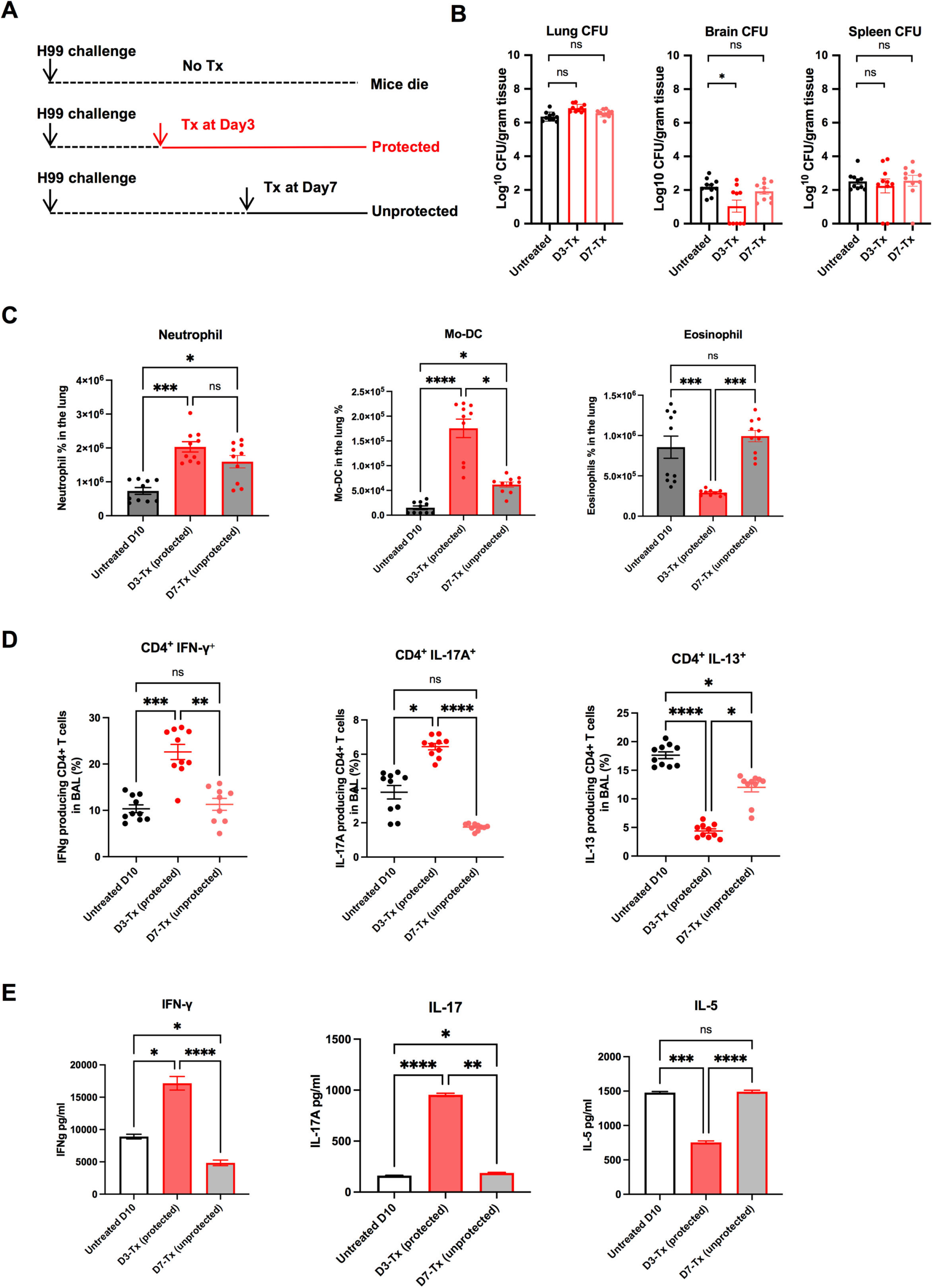
High dose and early intervention are required for the effectiveness of HK-fbp1 treatment against *Cryptococcus* infection. **(A)** Scheme of HK-fbp1 treatment strategy. WT C67BL/6 mice were infected with 1×10^4^ H99 at day 0. Mice were treated with 5×10^7^ HK-fbp1 on day3 (early treatment) or day7 (late treatment). **(B)** Fungal burden in the infected lungs, brains and spleens at day 10 post H99 infection from untreated mice, day 3 early treatment mice, and day 7 late treatment mice. Each symbol represents one mouse. Bars represent the mean ± the standard error of the mean. “*”, P<0.05 (determined by Mann-Whitney test). **(C)** Cellular infiltration of the lungs was analyzed by flow cytometry. Each symbol represents one mouse, and the data are accumulative from two independent experiments. Bars represent the mean ± the standard error of the mean. Each cell population was identified as CD45^+^ DAPI-negative live leukocytes. Mo-DCs were gated as CD11b^+^ Ly6G^-^ Ly6C^hi^ CD11c^+^ MHCII^+^, eosinophils were gated as CD11c^low^ SiglecF^+^, and neutrophil were CD11b^+^ Ly6C^int^ Ly6G^hi^. **(D)** Plots of cytokine production in CD4^+^ T cells gated as Thy1.2^+^ CD4^+^ CD8^-^ T cells in BALF. The frequencies of IFN-γ-, IL-17A-, and IL-13-producing CD4^+^ T cells in BALF were analyzed as shown. **(E)** CD4^+^ T cells were isolated from the lung-draining lymph nodes of untreated, day3 treated, day7 treated mice. Cytokine secretion in the presence of *Cryptococcus* antigen was examined by ELISA as described in Materials and Methods. The data shown are cumulative from two independent experiments with five mice per group and are depicted as the mean the mean ± the standard error of the mean. “****”, p < 0.0001; “***”, p < 0.001; “**”, p < 0.01; “*”, p < 0.05 (determined by One-way ANOVA nonparametric test for multiple comparisons).

We hypothesized that early treatment of HK-fbp1 likely shapes H99 immunogenicity and affects the development of protective immune response in the host, therefore preventing fungal dissemination in an effective manner. To test this hypothesis, we analyzed the recruitment of immune cells to the lung at day 10 post H99 infection. Our results showed that early treatment of HK-fbp1 induced an enhanced activation of innate immune response compared to the late treatment (Fig. 4C). We observed increased numbers of neutrophils and enhanced monocyte differentiation into monocyte-derived dendritic cells (Mo-DCs) in day 3-treated mice compared to the day 7-treated mice (Fig. 4C). Moreover, we overserved a decreased number of eosinophils in day 3-treated mice (Fig. 4C). Eosinophilia is a hallmark of Th2-dominated responses (24, 25), thus reduction in eosinophil numbers are indicative of reduced Th2 responses in Hk-fbp1-treated mice.

Previous studies have shown that Th1 (characterized by production of IFN-γ) and Th17 (characterized by production of IL-17) CD4^+^ T cells are important in defense of *Cryptococcus* infection in mouse models (9, 26). In contrast, Th2 responses (characterized by the production of IL-13) are harmful to the host during cryptococcosis (27–29). In our previous study we determine that monocytes and Mo-DCs are required for protection against *fbp1*Δ infection at least in part via activation of Th1 cells (20). Thus, we hypothesize that the enhanced Mo-DC maturation seen in in mice given HK-fbp1 at day 3 after H99 infection (Fig. 4C) help promote a protective T cell response. Therefore, we examined the cytokine profile of CD4^+^ T cells in the airway at day 10 post infection. We observed that early treatment with HK-fbp1 induced enhanced differentiation of IFN-γ-producing Th1 cells and IL-17A-producing Th17 cells in the airway (Fig. 4D) This increased protective response was accompanied by reduced differentiation of IL-13-producing Th2 cells (Fig. 4D). Reduced Th2 responses in HK-fbp1 treated mice are also indicated by reduced eosinophilia (Fig. 4C). Our aggregate observations suggest that early treatment with HK-fbp1 was able to shape CD4^+^ T cell polarization towards Th1 and Th17 responses and diminished detrimental Th2 differentiation. Similarly, *Cryptococcus*-specific CD4^+^ T cell responses measured in mLNs of mice treated early with HK-fbp1 also produced higher amounts of IFN-γ and IL-17A and lower amounts of IL-5 after *ex vivo* restimulation as compared to untreated controls or mice treated at day 7 (Fig. 4E). Collectively, these findings indicate that early treatment with HK-fbp1 induced enhanced recruitment of innate immune cells, as well as increased induction of Th1 and Th17 responses, and a lower Th2 response. These protective host immune responses help promote the containment of fungal cells in the lung and inhibition of fungal dissemination.

## Discussion

In this study we set out to examine the potential utility of HK-fbp1 as a therapeutic agent to treat cryptococcosis. The rationale for the studies presented stems from our earlier work where we demonstrated that vaccination with HK-fbp1 can provide potent protection against *Cryptococcus* infection by inducing a protective T cell response (20, 21). Our results showed that indeed, the HK-fbp1 vaccine is effective in treating early-stage *Cryptococcus* infection. However, the therapeutic value is limited when used to treat animals with disseminated cryptococcosis, indicating the T cell mediated immune response can contain fungal infection from dissemination before it happens.

To understand the role of T cells in HK-fbp1 mediated host protection, we performed CD4^+^ T cell depletion experiment in a previous vaccination study and found that the lack of CD4^+^ T cells triggered increased CD8^+^ cell expansion to provide protection (21). In this study, we depleted CD8^+^ T cells and both CD4^+^ and CD8^+^ T cells. While depletion of CD8^+^ T cells did not compromise the protection, we found that the animals depleted of both CD4^+^ and CD8^+^ T cells were no longer protected by the HK-fbp1 vaccine, suggesting that at least one T cell subset is needed to mount effective host protection. This result is consistent with the compensatory role of CD4^+^ and CD8^+^ T cells seen in other models of vaccine protection (30, 31), and is also consistent with a recent report in the *sgl1*Δ vaccination study (14). In that report, Dr. Del Poeta group also found that the presence of either CD4^+^ T cell or CD8^+^ T cells is sufficient to confer protection from the live *sgl1*Δ cell-based vaccine against *C. neoformans* challenge, but not in the host lacking both CD4^+^ and CD8^+^ T cells. A similar conclusion was also reported in the model of H99γ-based vaccine-induced protection (32). All these studies demonstrated the critical role of T cell mediated immunity in protection against *Cryptococcus* infection. Thus, our observations are consistent with findings in other models of vaccination.

Given the importance of CD4^+^ and CD8^+^ T cells in host defense against *C. neoformans* infection, we investigated the potential of utilizing HK-fbp1 vaccine as a therapeutic agent. We found that indeed, therapeutic administration of Hk-fbp1 can confer potent protection in mice treated early after fungal infection. The observation that the effective therapeutic potential is restricted to early administration is consistent with the interpretation that the mechanism of protection depends on the effective skewing of T cell differentiation towards protective Th1 and Th17 responses. In turn, protective immunity helps by blocking fungal dissemination. While we were testing the therapeutic value of our HK-fbp1 vaccine candidate, an independent study was reported using the *sgl1*Δ mutant based vaccine as a therapeutic agent, in which a similar treatment plan was used to treat mice infected by the parental *C. neoformans* strain (15). In that report, both heat-killed *sgl1*Δ cells and live *sgl1*Δ cells treatment of wild type infected mice significantly inhibited the *Cryptococcus* dissemination with reduced fungal burdens in the infected brains. Interesting, they also found that early treatment (day 3 post-infection) exhibited better outcome than later treatment (day 7 post-infection). The similar therapeutic outcome of these vaccine candidates in treating early stage cryptococcosis suggests they may share a very similar disease inhibition mechanism that is likely related to their ability to induce strong Th1 protective immunity although host immune responses were not examined in that study (15).

Previous studies suggest that infection with virulent strains of *C. neoformans* will induce a high Th2 response, which is detrimental to the host, and does not prevent disseminated infection (29, 33, 34). In this study we observed that mice treated with HK-fbp1 at day 3 after infection exhibited a high Th1 and Th17 protective responses, evident by increased production of IFN-γ and IL-17A cytokines and reduced Th2 cytokines (IL-13). In contrast, untreated mice and those treated at day 7 after infection showed lower Th1 responses and robust Th2 responses (IL5 and IL-13 producing T cells and increased eosinophilia). We conclude that administration of HK-fbp1 at day 7 after infection is unable to significantly alter the fate of T cells that already committed to a Th2 differentiation program.

Altogether, our study demonstrates that the HK-fbp1 vaccine candidate can not only be utilized as a prophylactic vaccine candidate to prevent multiple important invasive fungal infections, but it is also effective to treat early-stage *Cryptococcus* infection as a therapeutic agent. Although the utility of HK-fbp1 as a therapeutic agent may be restricted by the time of treatment and host conditions, further study and product improvement may lead to a more effective agent. In addition, it might be possible to develop combinational therapy where sequential of co-administration of HK-fbp1 vaccine and antifungal drugs may lead to improved control of fungal burden and better treatment outcomes.

## Acknowledgements

We thank Drs. Karthikeyan Nattarayan from Xue lab and Vanessa Espinosa from Rivera lab for technique support on this study. This study is supported by NIH grant R01AI141368 to AR and CX, and the Rutgers HealthAdvance Fund (partially supported through NIH U01HL150852) to CX. Studies in the Xue lab are also supported by NIH grants R01AI123315 and R21AI154318. AR holds an Investigators in the Pathogenesis of Infectious Disease Award from the Burroughs Wellcome Fund.

## Figure legends

**Figure S1.**
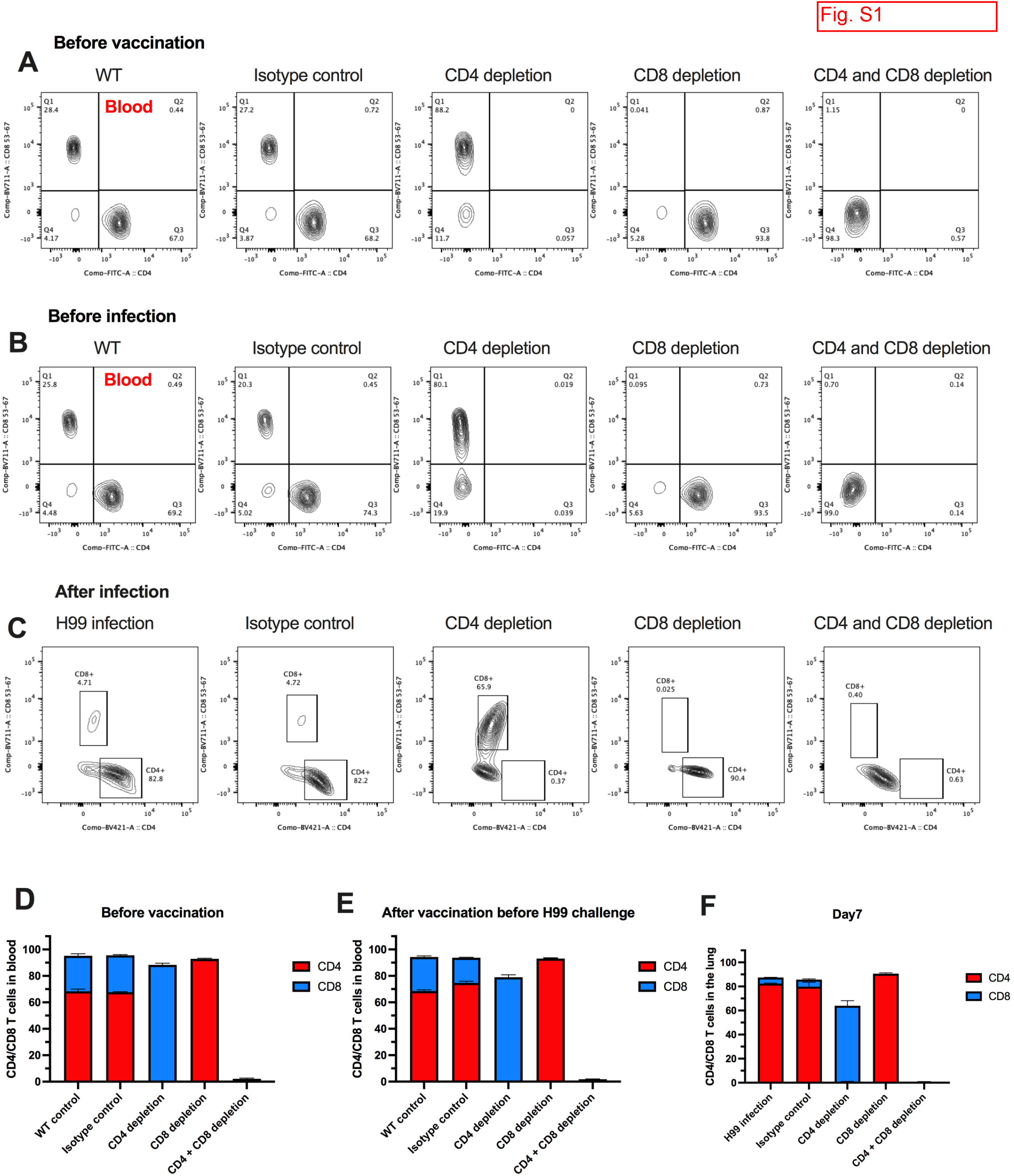
**(A and B)** Detection of CD4^+^ and CD8^+^ T cells by flow cytometry prior to first vaccination (day −43) **(A)** and prior to challenge infection (day −1) **(B)** in mice injected with either CD4 antibody, CD8 antibody or isotype antibody. Representative FACS plots of CD4^+^ and CD8^+^ T cells in blood samples of CD4^+^ T cells depleted mice, CD8^+^ T cells depleted mice, double depleted mice, isotype control treated mice and wild type infection only mice were shown. **(C)** Efficient depletion of CD4^+^ and CD8^+^ T cells in mice was confirmed by flow cytometry at day 7 post-challenge. Representative FACS plots of CD4^+^ and CD8^+^ T cells from lung tissues of CD4^+^ T cells depleted mice, CD8^+^ T cells depleted mice, double depleted mice, isotype control mice and wildtype infection only mice were shown. Each cell population was identified as CD45^+^, 4’,6-diamidino-2-phenylindole (DAPI)-negative live leukocytes. CD4^+^ T cells was gate as CD11b^-^, CD4^+^, CD8^-^; and CD8^+^ T cells were gated as CD11b^-^, CD4^-^, CD8^+^. **(D-F)** Percentages of CD4^+^ and CD8^+^ T cell in mouse blood at different times in the course of vaccination study: before vaccination **(D)**, before H99 challenge **(E)**, and at day 7 post-challenge **(F)**.

**Figure S2.**
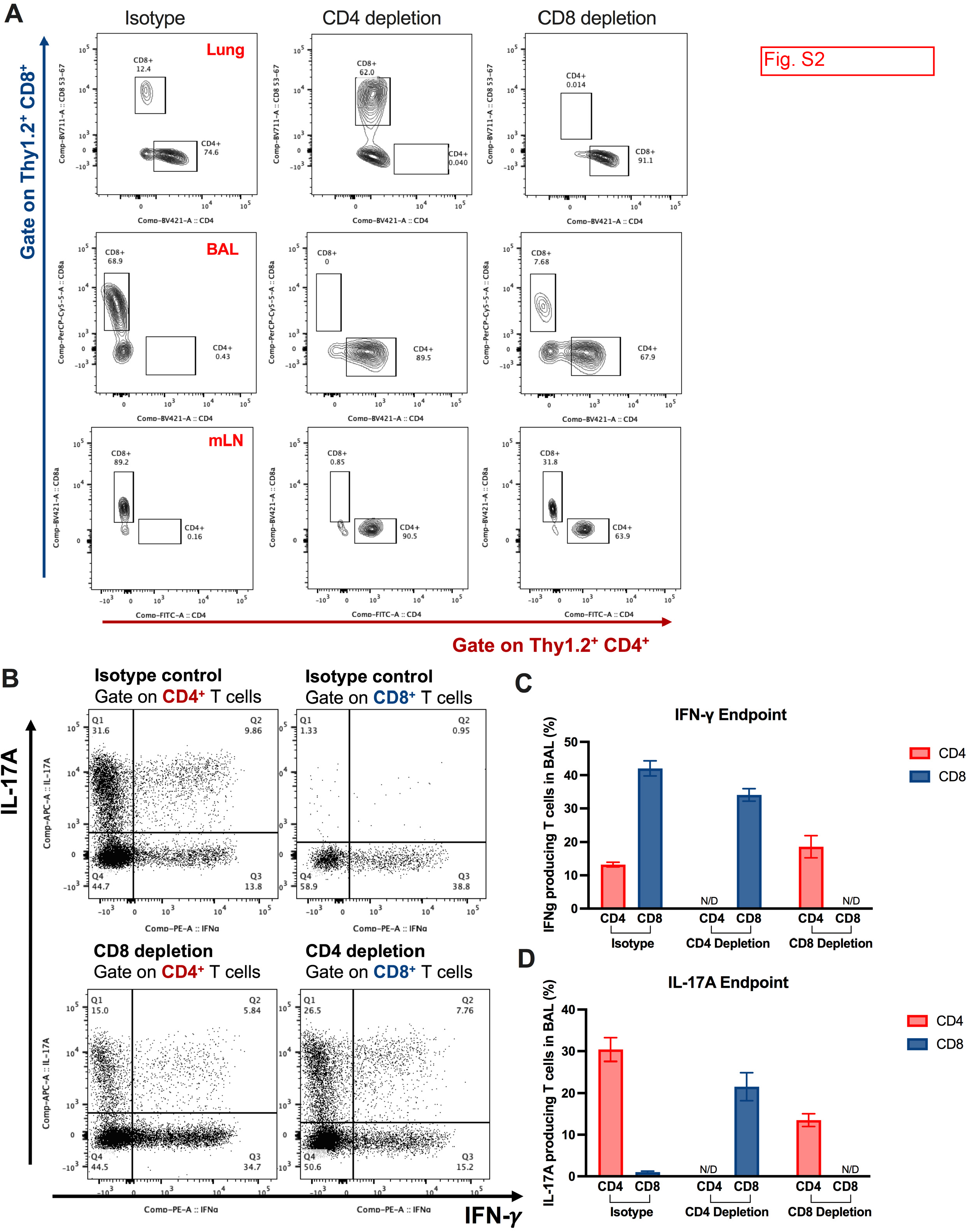
**(A)** At the endpoint of experiment (day 70), representative FACS plots of CD4^+^ T cells and CD8^+^ T cells in the lung, BALF, lung-draining lymph nodes in isotype control mice (left), CD4^+^ T cells depleted mice (middle), and CD8^+^ T cells depleted mice (right). **(B)** Representative FACS plots of cytokine production gate on CD4^+^ T cells or CD8^+^ T cells in isotype control mice (top), CD4 depleted mice (bottom), and CD8 depleted mice (bottom) at endpoint. **(C and D)** Cytokine expression at endpoint analyzed by ICCS. Plots of cytokine production in CD4^+^ T cells gated as Thy1.2^+^ CD4^+^ CD8^-^ T cells. Plots of cytokine production in CD8^+^ T cells gated as Thy1.2^+^ CD4^-^ CD8^+^ T cells. The frequencies of IFN-γ **(C)**, IL-17A **(D)** producing CD4^+^ or CD8^+^ in BALF were analyzed as shown in panel.

## References

1. Bongomin F, Gago S, Oladele RO, Denning DW. 2017. Global and Multi-National Prevalence of Fungal Diseases-Estimate Precision. J Fungi (Basel) 3.

2. Arastehfar A, Gabaldon T, Garcia-Rubio R, Jenks JD, Hoenigl M, Salzer HJF, Ilkit M, Lass-Florl C, Perlin DS. 2020. Drug-Resistant Fungi: An Emerging Challenge Threatening Our Limited Antifungal Armamentarium. Antibiotics (Basel) 9.

3. Perlin DS. 2011. Current perspectives on echinocandin class drugs. Future Microbiol 6:441–57.

4. Rivera A, Lodge J, Xue C. 2022. Harnessing the immune response to fungal pathogens for vaccine development. Annu Rev Microbiol 76:703–726.

5. Edwards JE, Jr., Schwartz MM, Schmidt CS, Sobel JD, Nyirjesy P, Schodel F, Marchus E, Lizakowski M, DeMontigny EA, Hoeg J, Holmberg T, Cooke MT, Hoover K, Edwards L, Jacobs M, Sussman S, Augenbraun M, Drusano M, Yeaman MR, Ibrahim AS, Filler SG, Hennessey JP, Jr. 2018. A fungal immunotherapeutic vaccine (NDV-3A) for treatment of recurrent vulvovaginal candidiasis-A Phase 2 randomized, doubleBlind, placebo-controlled trial. Clin Infect Dis 66:1928–1936.

6. Oliveira LVN, Wang R, Specht CA, Levitz SM. 2021. Vaccines for human fungal diseases: close but still a long way to go. NPJ Vaccines 6:33.

7. Caballero Van Dyke MC, Wormley FL, Jr. 2018. A call to arms: Quest for a Cryptococcal vaccine. Trends Microbiol 26:436–446.

8. Rajasingham R, Smith RM, Park BJ, Jarvis JN, Govender NP, Chiller TM, Denning DW, Loyse A, Boulware DR. 2017. Global burden of disease of HIV-associated cryptococcal meningitis: an updated analysis. Lancet Infect Dis 17:873–881.

9. Price MS, Perfect JR. 2011. Host defenses against cryptococcosis. Immunol Invest 40:786–808.

10. Wormley FL, Jr., Perfect JR, Steele C, Cox GM. 2007. Protection against cryptococcosis by using a murine gamma interferon-producing *Cryptococcus neoformans* strain. Infect Immun 75:1453–62.

11. Zhai B, Wozniak KL, Masso-Silva J, Upadhyay S, Hole C, Rivera A, Wormley FL, Jr., Lin X. 2015. Development of protective inflammation and cell-mediated immunity against *Cryptococcus neoformans* after exposure to hyphal mutants. MBio 6:e01433–15.

12. Upadhya R, Lam WC, Maybruck B, Specht CA, Levitz SM, Lodge JK. 2016. Induction of protective immunity to cryptococcal infection in mice by a heat-killed, chitosan-deficient strain of *Cryptococcus neoformans*. MBio 7:e01433–15.

13. Rella A, Mor V, Farnoud AM, Singh A, Shamseddine AA, Ivanova E, Carpino N, Montagna MT, Luberto C, Del Poeta M. 2015. Role of Sterylglucosidase 1 (Sgl1) on the pathogenicity of *Cryptococcus neoformans:* potential applications for vaccine development. Front Microbiol 6:836.

14. Normile TG, Rella A, Del Poeta M. 2021. *Cryptococcus neoformans* Deltasgl1 vaccination requires either CD4(+) or CD8(+) T cells for complete host protection. Front Cell Infect Microbiol 11:739027.

15. Normiie TG, Del Poeta M. 2022. Three models of vaccination strategies against cryptococcosis in immunocompromised hosts using heat-Killed *Cryptococcus neoformans* Deltasgl1. Front Immunol 13:868523.

16. Specht CA, Lee CK, Huang H, Tipper DJ, Shen ZT, Lodge JK, Leszyk J, Ostroff GR, Levitz SM. 2015. Protection against experimental cryptococcosis following vaccination with glucan particles containing *Cryptococcus* alkaline extracts. MBio 6:e01905–15.

17. Specht CA, Homan EJ, Lee CK, Mou Z, Gomez CL, Hester MM, Abraham A, Rus F, Ostroff GR, Levitz SM. 2022. Protection of mice against experimental cryptococcosis by synthesized peptides delivered in glucan particles. mBio 13:e0336721.

18. Liu TB, Wang Y, Stukes S, Chen Q, Casadevall A, Xue C. 2011. The F-Box protein Fbp1 regulates sexual reproduction and virulence in *Cryptococcus neoformans*. Eukaryot Cell 10:791–802.

19. Liu TB, Xue C. 2014. Fbp1-mediated ubiquitin-proteasome pathway controls *Cryptococcus neoformans* virulence by regulating fungal intracellular growth in macrophages. Infect Immun 82:557–68.

20. Masso-Silva J, Espinosa V, Liu TB, Wang Y, Xue C, Rivera A. 2018. The F-Box protein FFp1 shapes the immunogenic potential of *Cryptococcus neoformans*. MBio 9:e01828–17.

21. Wang Y, Wang K, Masso-Silva JA, Rivera A, Xue C. 2019. A heat-killed *Cryptococcus* mutant strain induces host protection against multiple invasive mycoses in a murine vaccine model. mBio 10:e02145–19.

22. Rivera A, Hohl TM, Collins N, Leiner I, Gallegos A, Saijo S, Coward JW, Iwakura Y, Pamer EG. 2011. Dectin-1 diversifies *Aspergillus fumigatus-specific* T cell responses by inhibiting T helper type 1 CD4 T cell differentiation. J Exp Med 208:369–81.

23. Sela M, Hilleman MR. 2004. Therapeutic vaccines: realities of today and hopes for tomorrow. Proc Natl Acad Sci U S A 101 Suppl 2:14559.

24. Leon B, Ballesteros-Tato A. 2021. cModulating Th2 Cell immunity for the treatment of asthma. Front Immunol 12:637948.

25. Jacobsen EA, Zellner KR, Colbert D, Lee NA, Lee JJ. 2011. Eosinophils regulate dendritic cells and Th2 pulmonary immune responses following allergen provocation. J Immunol 187:6059–68.

26. Rohatgi S, Pirofski LA. 2015. Host immunity to *Cryptococcus neoformans*. Future Microbiol 10:565–81.

27. Wiesner DL, Smith KD, Kashem SW, Bohjanen PR, Nielsen K. 2017. Different lymphocyte populations direct dichotomous eosinophil or neutrophil responses to pulmonary *Cryptococcus* infection. J Immunol 198:1627–1637.

28. Heung LJ. 2017. Innate immune responses to *Cryptococcus*. J Fungi (Basel) 3.

29. Holmer SM, Evans KS, Asfaw YG, Saini D, Schell WA, Ledford JG, Frothingham R, Wright JR, Sempowski GD, Perfect JR. 2014. Impact of surfactant protein D, interleukin-5, and eosinophilia on Cryptococcosis. Infect Immun 82:683–93.

30. Nanjappa SG, Heninger E, Wuthrich M, Sullivan T, Klein B. 2012. Protective antifungal memory CD8(+) T cells are maintained in the absence of CD4(+) T cell help and cognate antigen in mice. J Clin Invest 122:987–99.

31. Nanjappa SG, Klein BS. 2014. Vaccine immunity against fungal infections. Curr Opin Immunol 28:27–33.

32. Wozniak KL, Young ML, Wormley FL, Jr. 2011. Protective immunity against experimental pulmonary cryptococcosis in T cell-depleted mice. Clin Vaccine Immunol 18:717–23.

33. Goldman DL, Davis J, Bommarito F, Shao X, Casadevall A. 2006. Enhanced allergic inflammation and airway responsiveness in rats with chronic *Cryptococcus neoformans* infection: potential role for fungal pulmonary infection in the pathogenesis of asthma. J Infect Dis 193:1178–86.

34. Jain AV, Zhang Y, Fields WB, McNamara DA, Choe MY, Chen GH, Erb-Downward J, Osterholzer JJ, Toews GB, Huffnagle GB, Olszewski MA. 2009. Th2 but not Th1 immune bias results in altered lung functions in a murine model of pulmonary *Cryptococcus neoformans* infection. Infect Immun 77:5389–99.

